# *Mycobacterium tuberculosis* senses host Interferon-γ via the membrane protein MmpL10

**DOI:** 10.1101/2021.11.12.468344

**Authors:** Mohamed Ahmed, Jared Mackenzie, Robert Krause, Barry Truebody, Liku Tezera, Diana Garay-Baquero, Andres Vallejo, Katya Govender, John Adamson, Paul Elkington, Adrie JC Steyn, Alasdair Leslie

## Abstract

*Mycobacterium tuberculosis (Mtb)* is one of the most successful human pathogens and remains a leading cause of death from infectious disease. Interferon-γ (IFN-γ) is a central regulator of the immune defense against *Mtb*. Several cytokines have been shown to increase virulence of other bacterial pathogens, leading us to investigate whether IFN-γ has a direct effect on *Mtb*. We found that both recombinant and T-cell derived IFN-γ rapidly induced a dose-dependent increase in the oxygen consumption rate (OCR) of *Mtb*, consistent with increased bacterial respiration. This was also observed in clinical strains, but not in the vaccine strain Bacillus Calmette–Guérin (BCG), and did not occur for other cytokines tested, including TNF-α. IFN-γ binds to the cell surface of intact *Mtb*, but not BCG, whilst TNF-α binds to neither. Mass spectrometry analysis identified mycobacterial membrane protein large 10 (MmpL10) as the transmembrane binding partner. Consistent with this, IFN-γ binding and the OCR response was absent in a *Mtb Δmmpl10* strain and restored by complementation of the mutant strain. RNA-sequencing of IFN-γ exposed *Mtb* revealed a distinct transcriptional profile, including genes involved in virulence and cholesterol catabolism. Finally, exposure of *Mtb* cells to IFN-γ resulted in sterilization of bacilli treated with isoniazid (INH), indicating clearance of phenotypically resistant bacteria that persist in the presence of INH alone. Our data suggest a novel mechanism allowing *Mtb* to respond to host immune activation that may be important in the immunopathogenesis of TB and have use in novel eradication strategies.

**One-Sentence Summary:** IFN-γ is a critical component of effective immune defense in human tuberculosis yet its causative agent, *Mycobacterium tuberculosis* it able to sense this cytokine and increase virulence and respiration in response.

## Main Text

Tuberculosis (TB) remains a major public health challenge globally and is the leading cause of death from a single infectious disease globally. Infection with the causative pathogen, *Mycobacterium tuberculosis* (*Mtb*), initiates a series of highly complex inflammatory events, primarily in the lung, whose intricacy is not yet fully elucidated (*1*). The ancient relationship between the human host and *Mtb* has resulted in the acquisition of various mechanisms by *Mtb* to avoid immune destruction and establish persistent infection (*2*). Approximately 90% of infected individuals do not progress to clinical disease and, although the determinants of protective immunity against *Mtb* infection are not fully understood, several indispensable factors have been identified. The importance of adaptive immunity in humans, for example, is indicated by a direct correlation between the loss of CD4 T cells due to HIV infection and increasing risk of developing active TB. In addition, the importance of proinflammatory cytokines such as tumor necrosis factor-α (TNF-α), interleukin-1β (IL-1β), interleukin-12 (IL-12) and interferon-γ (IFN-γ) is demonstrated by the association between genetic deficiencies in their signaling pathways and increased risk of TB disease (*3*). IFN-γ is believed to be a particularly important component of the adaptive response to *Mtb* infection as activation of macrophages by IFN-γ is required to restrict mycobacterial replication (*4, 5*).

In contrast to these observations, overexpression of IFN-γ by CD4 T cells causes mice to succumb rapidly to *Mtb*, which is prevented by expression of the inhibitory receptor programmed cell death protein (PD-1) (*6*). Blockade of the PD-1 axis in cancer patients, which re-activates IFN-γ production by T cells, has led to numerous instances of latent TB reactivation (*7*). How inhibition of this pathway causes TB reactivation is, however, not well understood. Intriguingly, several bacterial pathogens directly sense and respond to host cytokines in order to facilitate their survival and transmission. For example, virulent *Escherichia coli* (*E. coli*), accelerates growth in response to IL-1β, but avirulent *E. coli* does not, suggesting that detection of host immune signals is an adaptation in pathogenic strains (*8*). Similarly, Gram-negative bacteria have been found to interact with TNF-α, leading to an increase in cellular invasion (*9*), whilst sensing of human IFN-γ drives *Pseudomonas aeruginosa* (*P. aeruginosa*) towards a virulent phenotype (*10*). In light of these observations and others, we hypothesize that *Mtb* directly senses IFN-γ as part of a countermeasure to host immunity.

To determine if there was any direct physiological effect of human IFN-γ on *Mtb*, we made use of the Agilent Seahorse XF Analyzer, a highly sensitive analytical platform capable of measuring changes in bacterial respiration (*11*). Upon addition of recombinant human IFN-γ, *Mtb* H37Rv rapidly increased in oxygen consumption rate (OCR) in a dose-dependent manner (Fig. 1A). By comparison, recombinant human TNF-α had no effect on *Mtb* OCR (Fig. 1B). To rule out the possibility that this observation was due to the use of recombinant human IFN-γ, we stimulated T cells non-specifically for 48 hours and then used the culture supernatant to stimulate *Mtb*. This also triggered a rapid increase in *Mtb* OCR, which was abrogated by depleting IFN-γ using a monoclonal antibody (Fig. 1C) and confirmed that the effect was specific to IFN-γ. These data also suggest that *Mtb* can sense physiologically relevant concentrations of IFN-γ. Likewise, no effect on *Mtb* OCR was observed when the recombinant cytokines IL-6, IL-1β, GM-CSF, M-CSF, IL-4 and IL-10 (Fig. 1D) were used. We also confirmed that recombinant murine IFN-γ had the same effect on *Mtb* OCR, implying that this phenotype may be relevant in studies using mouse models of TB (Fig. S1). To further validate this phenotype, we next tested two clinical *Mtb* isolates, derived from human TB subjects recruited from TB clinics in the Durban area and passaged only twice. We confirmed that both strains upregulated the OCR in response to IFN-γ (Fig. 1F). In sum, these data suggest that *Mtb* cells directly interact with IFN-γ, causing an increase in bacillary respiration.

**Figure 1:**
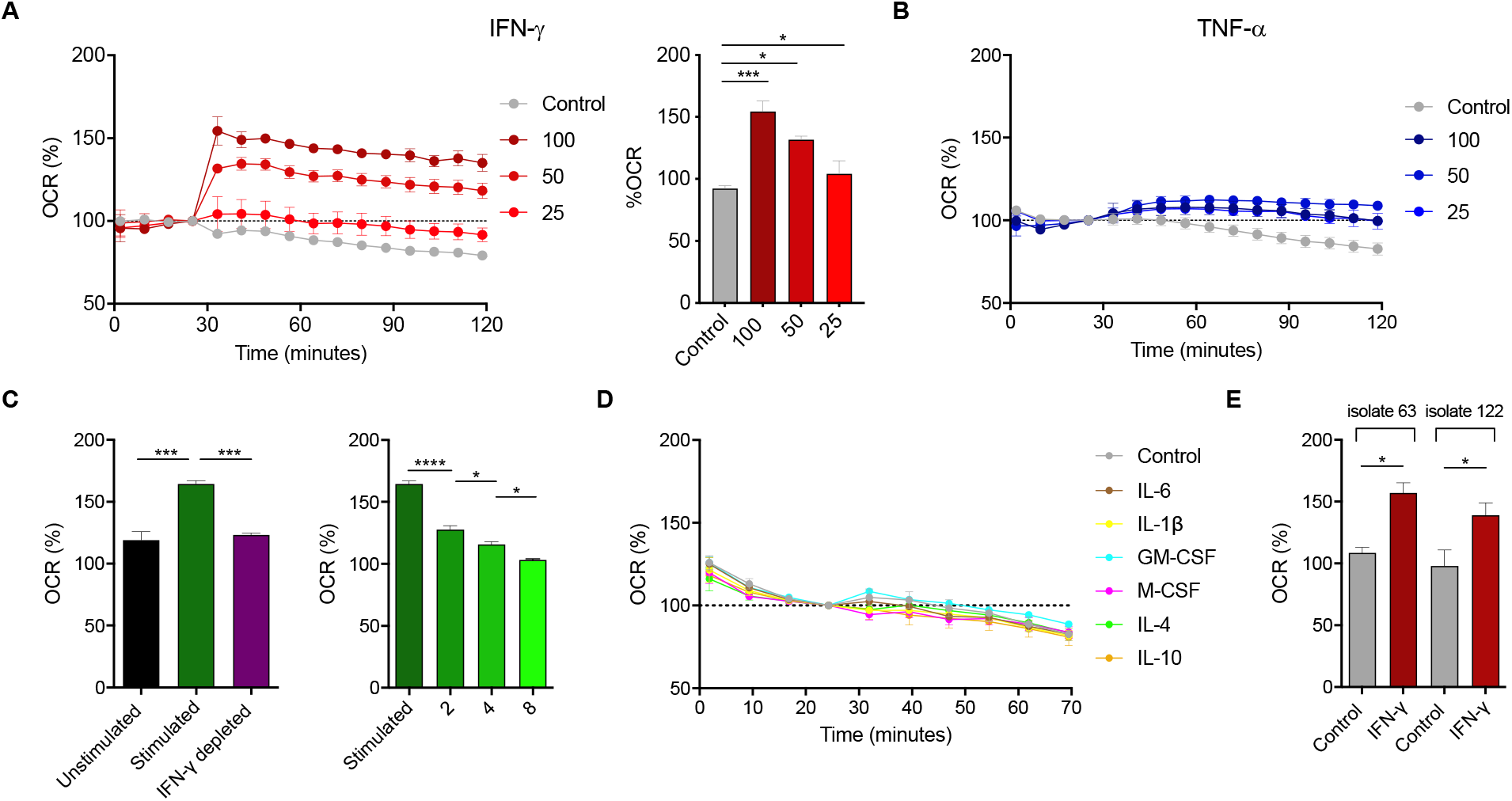
IFN-γ stimulates respiration in *Mtb*. Addition of recombinant human IFN-γ to *Mtb* at indicated concentrations (ng/mL) induces dose-dependent increase in OCR (A) whilst no effect was observed with recombinant human TNF-α (B). Media derived from stimulated (anti-CD3/CD28 treated) PBMC media, likewise increased OCR, which was abrogated upon IFN-γ depletion (C). Dilutions of stimulated media led to a dose-response reduction of the effect. This effect appears to be specific to IFN-γ as several cytokines failed to induce increased respiration (D). Clinical *Mtb* isolates also increased OCR in response to recombinant human IFN-γ (E). Data are shown as mean ± SEM and represent at a minimum two independent experiments. Tukey’s correction multiple-comparison test was used for the statistical analysis.

Following the observation that IFN-γ induced an increase in *Mtb* respiration, we reasoned that a physical interaction must occur. The outer membrane of *Mtb* contains various proteins that facilitate transport of molecules, indicating that *Mtb* has evolved mechanisms for binding macromolecules via surface components (*12*). To test whether *Mtb* can interact with IFN-γ, we incubated formalin-fixed whole *Mtb* with recombinant IFN-γ followed by staining with a fluorescent anti-IFN-γ antibody. Subsequent flow cytometry analysis indicates that IFN-γ binds to the surface of *Mtb* in a dose dependent manner (Fig. 2A). This result was confirmed by enzyme-linked immunosorbent assay (ELISA) (Fig. S2), and by confocal microscopy (Fig. 2C). In contrast, TNF-α, which failed to induce an increase in OCR, did not bind to *Mtb* using the same experimental setup (Fig. 2B). Interestingly, IFN-γ did not bind to the attenuated vaccine strain, Bacillus Calmette–Guérin (BCG) (Fig. 2D). In addition, no change in OCR was detected following addition of IFN-γ (Fig. 2E). Taken together, these data indicate that the ability of pathogenic mycobacteria to respond to IFN-γ stimulation is linked to a direct interaction with the cytokine at the cell surface.

**Figure 2:**
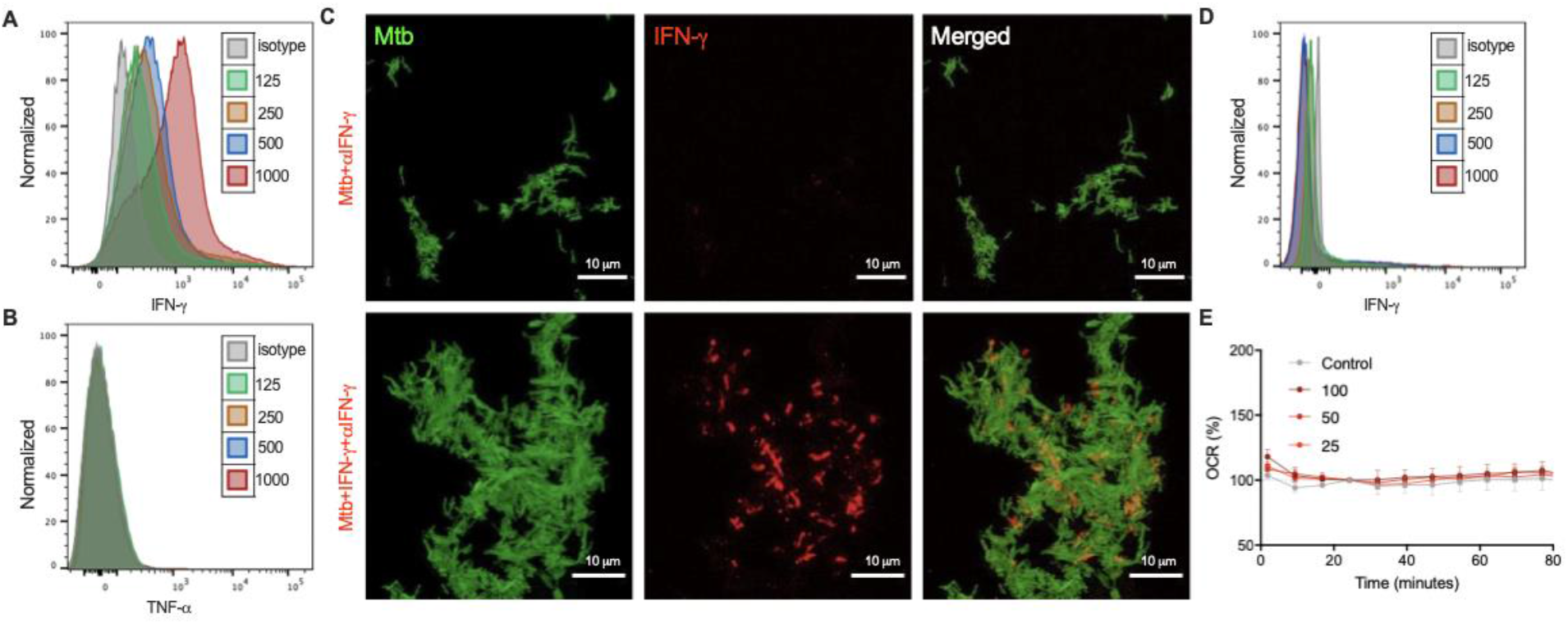
IFN-γ binds to surface of *Mtb* but not BCG. Flow cytometry shows that recombinant human IFN-γ (A) but not recombinant human TNF-α (B) binds to *Mtb* in a dose-dependent manner at the indicated concentrations (ng/mL). Confocal microscopy confirmed binding of recombinant human IFN-γ to individual *Mtb*-GFP (B). Top panel shows control sample; bottom panel shows fully stained sample. In contrast to *Mtb*, BCG does not bind IFN-γ (C) nor increase respiration in response to IFN-γ (D). Data are shown as mean ± SEM and represent at a minimum two independent experiments. Tukey’s correction multiple-comparison test was used for the statistical analysis.

We next sought to identify the binding partner of IFN-γ using a proteomics approach (Fig. 3A). Whole cell lysate of *Mtb* was separated by non-denaturing gel electrophoresis, transferred to polyvinylidene difluoride (PVDF) membrane by Western blotting and incubated with IFN-γ. The PVDF membranes were then stained with anti-IFN-γ antibody, revealing a distinct IFN-γ binding band (Fig. 3B). The corresponding area was excised from the reference gel and analyzed by LC/MS-MS mass spectrometry. Multiple experiments yielded several peptides corresponding to a number of different *Mtb* proteins (Table S1). Considering that IFN-γ binds on the surface of *Mtb* we narrowed this down to proteins located within the bacterial membrane, leaving only a single candidate, mycobacterium membrane protein large 10 (MmpL10) (Fig. 3C). The same unique peptide of MmpL10 was consistently observed in three independent experiments (Fig. S3), strongly suggesting an interaction between MmpL10 and IFN-γ. To test this putative IFN-γ binding partner, we obtained a *Δmmpl10* transposon mutant of *Mtb* (*13*) and assessed binding to IFN-γ as above. The *Δmmpl10* mutant did not bind IFN-γ (Fig. 3D), and complementation of the *Δmmpl10* mutant with wildtype *mmpl10* (*rv1183*) restored the binding of IFN-γ (Fig. 3E). In addition, the *Δmmpl10* mutant did not increase the OCR in response to IFN-γ, but this was restored in the *mmpl10*-complement (Fig. 3E). Indeed, overexpression of *mmpl10* in the complemented strain relative to *Mtb* H37Rv, as evidenced by a higher median fluorescent intensity in the flow cytometric binding experiment, was matched by a greater increase in OCR. To exclude the possibility that non-specific disruption of the bacterial membrane through mutation of a transmembrane protein affected the ability of *Mtb* to respond to IFN-γ, we tested the effect of IFN-γ on additional *Δmmpl* mutants available (MmpL-4, 5, 8 and 11), all of which are predicted transmembrane proteins. These analyses confirmed that only *mmpl10* mutation abrogated the metabolic response to IFN-γ (Fig. S4; Table S2). Having identified the likely binding partner for IFN-γ in *Mtb* we analyzed the sequence of MmpL10 in BCG, which did not bind or respond to IFN-γ. Intriguingly, the *mmpl10* gene in BCG is 100% homologous to that in *Mtb* H37Rv. To explore this further, we tested whole bacterial lysates of both BCG and the *Δmmpl10* mutant used above and found the same distinct IFN-γ binding band in both as observed in *Mtb* H37Rv (Fig. 3F). These data suggest that the failure of *Δmmpl10* and BCG to bind and respond to IFN-γ is due to differences in the localization or accessibility of MmpL10 rather than changes in the protein structure.

**Figure 3:**
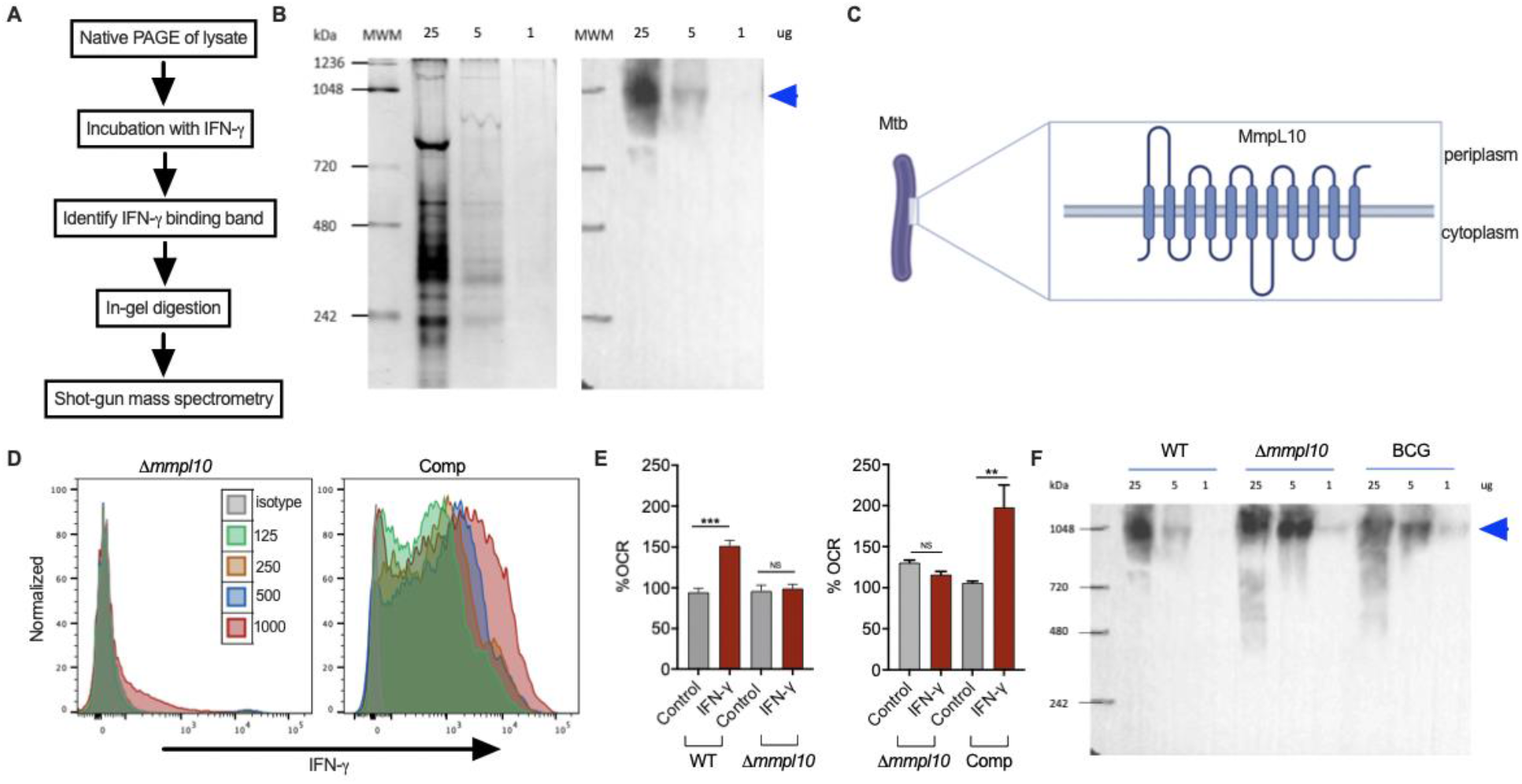
MmpL10 is the binding partner of IFN-γ. A proteomics approach to identify proteins in *Mtb* lysate that bind IFN-γ (A). Proteins from native page of *Mtb* lysate (B, left panel) were transferred to PVDF membrane. Subsequent staining of the membrane yielded an immunoreactive band to IFN-γ (B, right panel). Subsequent analysis using mass spectrometry revealed the band to contain peptides corresponding to MmpL10 (C). Binding of IFN-γ to *Δmmpl10* was abolished, while it was restored in the *mmpl10*-complemented strain (D). Similarly, the effect on OCR in response to IFN-γ was abolished in *Δmmpl10* and restored in *mmpl10*-complemented strain (E). Immunoreactive bands were also observed in lysates of *Δmmpl10* and BCG (F). Data are shown as mean ± SEM and represent at a minimum two independent experiments. Tukey’s correction multiple-comparison test was used for the statistical analysis.

To investigate the potential downstream effects of IFN-γ sensing by *Mtb* we turned to a 3D tissue-like model that mimics several key aspects of the human TB granuloma. Consistent with earlier experiments using this system, we found that supplementation of IFN-γ in the culture media increased bacterial growth in a dose-dependent manner (Fig. 4A) (*14*). In addition, and not previously tested, depletion of IFN-γ in this model significantly reduced bacterial burden by day 14 in culture (Fig. 4B). To determine whether stimulation with IFN-γ was linked to a transcriptional program in *Mtb*, we incubated multiple separate *Mtb* cultures for 18 hours in the presence of the cytokine and sequenced the extracted RNA at mid-log phase growth. Paired-end reads were aligned to the H37Rv reference transcriptome and counts per gene were obtained. Principal component analysis showed a strong batch effect, indicating a high degree of variability between different cultures (Fig. 4C). Despite this, however, a separation of IFN-γ stimulated and unstimulated bacteria was observed, indicating the transcriptional reprogramming of *Mtb* in response to IFN-γ. Interquartile range analysis showed no outliers (Fig. S5). Bioinformatic analysis of differentially expressed genes revealed that treatment with IFN-γ caused the significant regulation of several genes (*p* value < 0.001, *q* value <0.2, Table S3), including upregulation of *vapC14* (rv1953) and *esxP* (rv2347c), encoding putative virulence factors (Fig, 4C). Recently, VapC4, which, like *vapC14* is an RNase toxin, was shown to activate stress survival pathways in *Mtb* (*15*). The Esx-1 secretion system is an exporter of virulence factors including the ESAT-6-like protein encoded in *esxP* (*16*). Whilst the function of *esxP* is not known, the protein was one of a limited number found to be differentially expressed by *Mtb* H37Rv compared to attenuated strain H37Ra, implying a potentially important role in bacterial virulence (*17*). In addition, the genes *fadD3* (rv3561) and *ipdF* (rv3559c), both of which are involved in cholesterol catabolism, were significantly downregulated (Fig. 4D) (*18, 19*). Cholesterol has been identified as an important carbon source for *Mtb* even in the presence of alternative nutrients (*20*); and degradation of cholesterol has shown to be necessary for intracellular growth in macrophages (*21*). Taken together, these data demonstrate that virulent *Mtb* has the ability to sense host IFN-γ, via the MmpL10 protein, leading to transcriptional changes that may enhance its virulence and survival.

**Figure 4:**
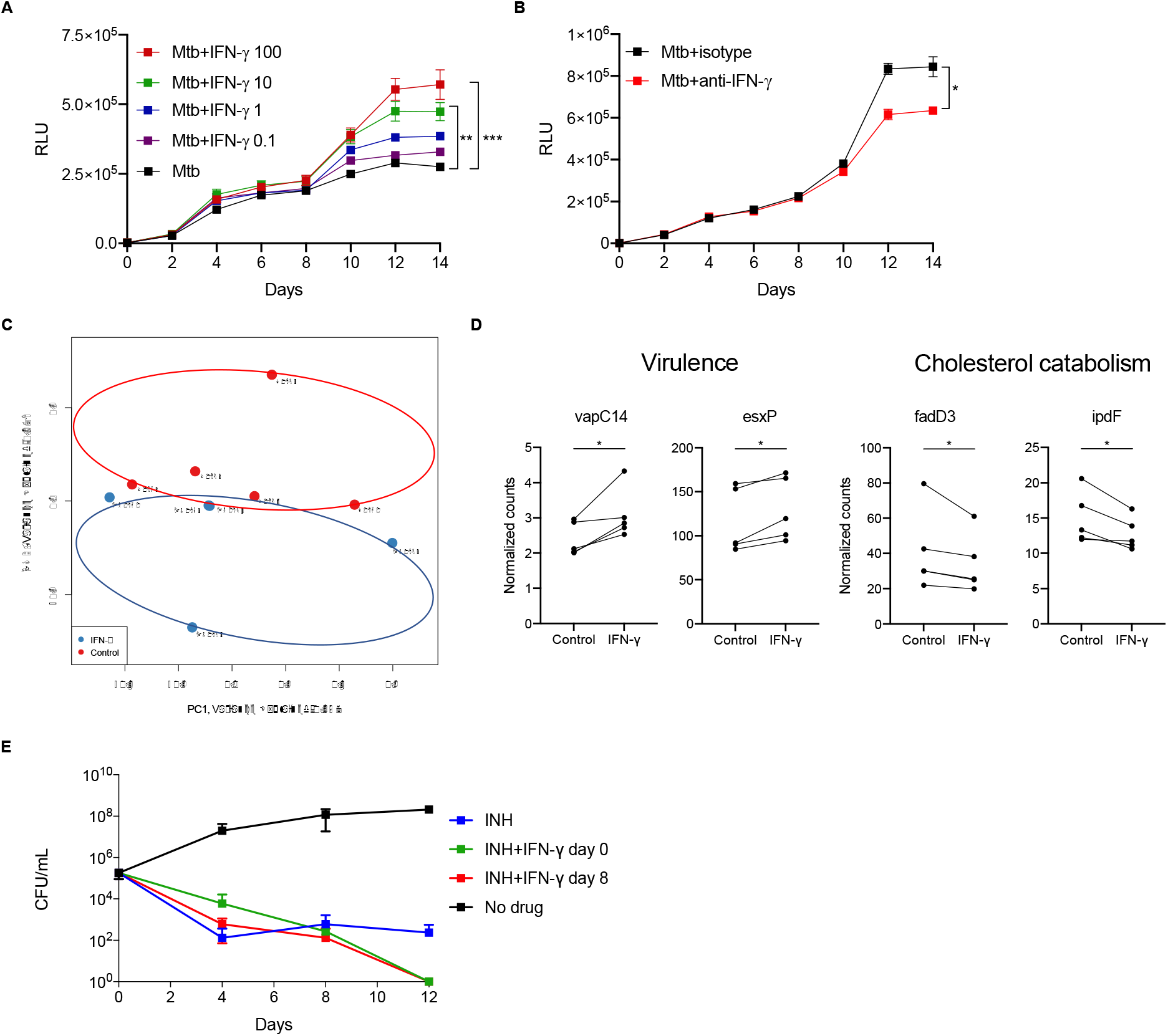
Distinct effects of IFN-γ on *Mtb*. IFN-γ increases bacterial growth (A) in a 3D granuloma model measured by relative light units (RLU) in a dose-dependent manner at the concentrations (ng/mL) indicated. Conversely, blockade of IFN-γ within the model reduces bacterial growth (B). A distinct transcriptional profile is observed in *Mtb* when exposed to IFN-γ when analyzed by RNAseq and PCA, (C) and IFN-γ regulates several genes of interest associated with virulence and cholesterol catabolism (D). IFN-γ enhances INH-mediated killing of *Mtb* in broth to sterilize cultures at day 12 (E). Data are shown as mean ± SEM and represent at a minimum two independent experiments. Tukey’s correction multiple-comparison test and Mann-Whitney test was used for the statistical analysis.

Finally, we sought an orthogonal method to confirm that *Mtb* upregulates its metabolism in response to IFN-γ stimulation. A major cause of antibiotic treatment failure is thought to be the existence of a population of persister bacteria. These are genetically susceptible organisms that are phenotypically resistant to certain antimicrobial drugs through a variety of mechanisms, including a reduction in metabolic activity (*22*). The efficacy of the pro-drug isoniazid (INH), for example, is dependent on its conversion by the catalase KatG into its active form. Therefore, sterilizing *Mtb* cultures with INH requires activation of persisters (*23*). We therefore hypothesized that IFN-γ stimulation of *Mtb* respiration should prevent the bacilli from entering a metabolically inactive, drug tolerant state. To test this, we cultured *Mtb* in the presence of INH, with or without the addition of IFN-γ. After 12 days, culturable bacteria were still detectable in the presence of INH alone, indicating the development of persisters as expected (Fig 4F). In contrast, the addition of IFN-γ, either at the start of the experiment, or on day 8, resulted in culture sterilization by day 12 (Fig. 4F). On its own, IFN-γ did not affect *Mtb* growth, ruling out the possibility of a direct toxic effect (Fig. S6) These findings strongly support the observation that exogenous IFN-γ stimulates *Mtb* metabolism, which prevents the formation of drug tolerant persisters, thereby enhancing killing by INH.

Here, we present evidence that *Mtb* possesses the ability to respond to host IFN-γ, thought to be a key mediator in the immune defense to this pathogen. We show that virulent *Mtb* binds to IFN-γ and exhibits a dose-dependent increase in metabolic activity, but not to other cytokines tested, including TNF-α. This is also true for clinical strains, but not for the avirulent vaccine strain BCG. We identify the transmembrane protein MmpL10 as the putative binding partner for IFN-γ, by showing that the ability to bind and respond to IFN-γ is lost in a *Δmmpl10* mutant strain and is restored by *mmpl10* complementation. In addition, RNA sequencing of *Mtb* demonstrates that IFN-γ induces a distinct transcriptional response, including the upregulation of virulence factors. Importantly, killing of *Mtb* by INH was enhanced by IFN-γ, consistent with its ability to induce bacterial respiration. Together, our findings indicate the existence of a novel mechanism that allows *Mtb* to adjust its metabolism by sensing a key host cytokine produced in the immune response to this pathogen. This suggests an evolutionary adaptation to the host immune response against this highly successful pathogen.

Adaptation to host immune mediators has been shown in other bacteria, lending biological plausibility to this observation. As a case in point, *P. aeruginosa* has been shown to sense host IFN-γ through the membrane porin OprF, resulting in expression of the virulence factor PA-I (*10*). Interestingly, increased IFN-γ does not enhance clearance of *P. aeruginosa*, which could be linked to the subversive action of OprF (*24*). As OprF and MmpL10 share little homology in structure or function this appears to be a shared adaption to similar immune pressures and not an example of horizontal gene transfer between bacterial pathogens. The transmembrane protein family of MmpLs are involved in the establishment of the mycobacterial cell envelope. MmpL10 specifically, is thought to be required for the translocation of diacyltrehaloses (DAT) across the plasma membrane, where they are further acylated to generate penta-acyltrehaloses (PAT) (*25*). Interestingly, *Mtb* mutants lacking the ability to synthesize DAT were equally infectious as wildtype *Mtb* in mice by either aerosol or intravenous infection, whilst *Δmmpl10* mutant was highly attenuated when given intravenously (*26, 27*). It is therefore possible that MmpL10 has virulence functions beyond glycolipid transport.

The phenomenon, known as “protein moonlighting”, of highly conserved proteins involved in metabolic regulation or the cell stress response having a range of additional biological actions which are involved in bacterial virulence, has been described in several bacteria including *Mtb* (*28*). Indeed, another member of the MmpL family, the integral membrane mycolic acid transporter MmpL3, was recently shown to have many binding interactions unrelated to its primary function that allow it to coordinate cell wall deposition during cell septation and elongation (*29*). The precise function and mode of action of transmembrane proteins in *Mtb* is complicated by the presence of the outer layers of mycobacteria, which separate the plasma membrane from the environment. However, the fact that the BCG lysate, in which MmpL10 is identical, binds IFN-γ but whole BCG does not, indicates that MmpL10 is oriented differently in pathogenic *Mtb*. Importantly, comparative analysis of purified membrane fractions of H37Rv and BCG by mass spectrometry revealed several proteins that were significantly more abundant in the H37Rv, despite being genetically identical, including MmpL10 (*30*).

Although IFN-γ is necessary for effective TB immunity, it is also clear that excessive Th1 responses, and IFN-γ in particular, can exacerbate disease. As highlighted above, a number of studies reported on the development of active TB disease in cancer patients receiving anti-PD-1 therapy, which enhanced IFN-γ production by *Mtb*-specific CD4 T cells (*31*). In an earlier work, it was shown that adolescents who exhibited the most intense reactions to tuberculin were more likely to develop TB as adults, many years after the initial tuberculin skin test (*32*). A recent meta-analysis of 34 longitudinal studies reporting the baseline magnitude of the IFN-γ response to *Mtb*, showed that higher levels IFN-γ are consistently associated with a greater risk of active TB (*33*). Supporting these clinical observations in humans, reports from animal models show that PD-1 blockade in macaques caused TB reactivation and PD-1 deficient mice have exaggerated IFN-γ responses and are highly sensitive to *Mtb* infection (*34, 35*). Interestingly, however, the control of BCG is enhanced in PD-1 deficient mice, where it is associated with a significant increase in antigen-specific IFN-γ production by CD4 T cells (*36*). Whilst there are many genetic reconfigurations in BCG that contribute to its attenuation, it is possible that the inability to detect and respond to host IFN-γ is a contributing factor. Likewise, although there are multiple potential reasons why excessive IFN-γ production could be detrimental to the host control of *Mtb*, the ability of the pathogen to respond to these conditions, however they arose, could contribute to the rapid disease progression observed.

The long period of human-*Mtb* co-evolution has left a genetic imprint manifesting itself in mechanisms of immune resistance and immune evasion in both host and pathogen. The advantage of host IFN-γ sensing is not clear from this current study, but the observation that treatment with IFN-γ leads to culture sterilization by INH, indicates that sensing this cytokine drives bacteria out of a persister state. *Mtb* can persist in immune competent hosts as a latent, or quiescent infection for years, which may represent an analogous “persister” state (*37*). It is speculated that the ability to develop into a latent infection may have evolved to facilitate spread when human populations were low, requiring the periodic breaking of latency for onward transmission (*38*). It is possible, therefore, that the ability to sense IFN-γ evolved as a mechanism to reactivate latent *Mtb* infection when the environment was favorable for transmission, for example during a respiratory infection. Considering that IFN-γ increases *Mtb* respiration, it may be possible to leverage this effect in several ways, for example to augment INH therapy, or to improve the detection of non-culturable bacteria in clinical samples (*39*). In this study we have demonstrated a novel effect of IFN-γ on *Mtb*, knowledge of which is critical to inform design of future TB therapies or vaccines to control the TB epidemic.

## Acknowledgments

We are grateful to James Millard (AHRI) for providing clinical isolates and Dr David Johnston of the Biomedical Imaging Unit (University of Southampton) for assistance with confocal imaging. Outr thanks to Amanda Ardain for proofreading the manuscript.

## Funding

MA

Sub-Saharan African Network for TB/HIV Research Excellence (SANTHE), PhD Fellowship

PE

Medical Research Council, MR/P023754/1

Medical Research Council, MR/N006631/1

AJCS

National Institutes of Health, R01AI134810

AL

Wellcome Trust, Senior Research Fellowship (210662/Z/18/Z)

Medical Research Council, MRC Global Challenges Research Fund (MR/P023754/1)

## Author contributions

Conceptualization: MA, JM, LT, PE, AJCS, AL

Methodology: MA, JM, RK, BT, LT, DGB, AV, KG, JA, PE, AJCS, AL

Investigation: MA, JM, RK, BT, LT, DGB, AV, KG, JA, PE, AJCS, AL

Visualization: MA, PE, AL

Funding acquisition: MA, PE, AJCS, AL

Project administration: AJCS, AL

Supervision: PE, AL

Writing – original draft: MA, PE, AL

Writing – review & editing: MA, LT, DBG, PE, AJCS, AL

## Data and materials availability

RNAseq dataset will be published on Gene Expression Omnibus (GEO). Programming code will be made available on Github

## Supplementary Materials

Materials and Methods

Supplementary Text

Figs. S1 to S6

Tables S1 to S3

References (40)

