## Supplemental data for "*Mycobacterium tuberculosis* senses host Interferon-γ via the membrane protein MmpL10"

**This PDF file includes:**

Materials and Methods
Figs. S1 to S6
Tables S1 to S3

### Materials and Methods

#### Bacterial strains and growth conditions

All mycobacterial strains were cultured in Middlebrook 7H9 broth (Difco) (supplemented with 10% OADC, 0.2% glycerol and 0.01% Tyloxapol (Sigma-Aldrich) at 37°C. *Mtb*-GFP was grown in media containing hygromycin 50 μg/mL. *Mtb Δmmpls* were obtained from the John's Hopkins University Mutant Library (*13*) and grown in media containing kanamycin 25 μg/mL; *Mtb Δmmpl10::*aph complement was grown in media containing 25 μg/mL kanamycin with 50 µg/mL hygromycin. Bioluminescent *Mtb* H37Rv was grown in media containing 25 μg/mL kanamycin. Live bacteria were used in all Seahorse experiments. Whole bacteria fixed in 4% paraformaldehyde (PFA) were used for Enzyme Linked immunosorbent assay (ELISA), confocal microscopy and flow cytometry experiments. Bacterial lysates were prepared for Western blotting. *Mtb* CFU counting was performed by serial dilution in PBS-Tween 80 (0.05%) (Sigma Aldrich) on Middlebrook 7H11 agar with OADC (Difco) unless stated otherwise.

#### *Mtb* OCR measurements

OCR measurements were conducted as described previously (*11*). Briefly, an XFe96 Extracellular Flux Analyser (Seahorse Biosciences) was used to measure the OCR of *Mtb* bacilli. These bacilli were adhered to the bottom of a Cell-Tak coated XF cell culture microplate (Seahorse Biosciences) at $2*10^6$ bacilli per well. Assays were carried out in unbuffered 7H9 media (pH 7.35) without a carbon source. In general, basal OCR was measured for 21 min before the automatic addition of the cytokines (Peprotech) or other stimulants through the drug ports of the sensor cartridge. All OCR figures indicate the point of each addition as a dotted line. OCR data points are representative of the average OCR during 4 min of continuous measurements in the transient microchamber, with the error being calculated from the OCR measurements taken from at least three replicate wells by the Wave Desktop 2.2 software (Seahorse Biosciences).

**IFN-γ binding by ELISA and flow cytometry**

Binding of IFN-γ to *Mtb* was assessed by mixing varying concentrations of IFN-γ with whole bacterial cells fixed in 4% PFA and incubating overnight at 4°C. Bacteria were pelleted and resuspended in 0.1% Tween in PBS before seeding onto microtiter wells for 2 hours at 37°C. After washing, Ultra-LEAF anti-human IFN-γ antibody (Biolegend, clone B27) was added to the wells which were incubated for a further 2 hours at room temperature. Anti-mouse HRP was then added for 1 hour at room temperature. TMB solution (Sigma Aldrich) was used as a substrate and 1 M sulfuric acid as stop solution. Optical density was read at 450 nm measured with GloMax Discover microplate reader (Promega). For flow cytometry, after overnight incubation with IFN-γ, bacterial pellets were stained with human anti-IFN-γ Brilliant Violet 421 (Biolegend, clone 4S.B3) or anti-TNF-α Alexa Fluor 700 (BD, clone MAb11) for 1 hour at room temperature. Data was acquired using BD Aria Fusion cytometer and analyzed using FlowJo Software v.10.

**Confocal microscopy**

*Mtb*-GFP fixed in 4% PFA was mixed with IFN-γ overnight at 4°C. Bacteria were stained with anti-human IFN-γ APC (Biolegend, clone 4S.B3) for 1 hour at room temperature. After washing, cells were resuspended in mountant and fixed to the slide. Samples were imaged using an Olympus IX81 microscope and images were exported as lif files and opened in ImageJ.

**Stimulation of PBMCs**

Peripheral blood mononuclear cells (PBMCs) were isolated from whole blood by centrifugation on Ficoll-Paque (Sigma), resuspended in RPMI 1640 medium (Thermo Fisher) containing 10% Fetal Calf serum, 1% ampicillin and seeded at $2*10^5$ cells/well in 96 well plates. Cells were cocultured with Dynabeads Human T-Activator CD3/CD28 (Invitrogen) to stimulate T cells at 37°C in 5% $CO_2$ for 48 hours. Supernatants were recovered by centrifugation. IFN-γ was depleted from the supernatant with Ultra-LEAF anti-human IFN-γ antibody (Biolegend, clone B27).

**Detection of IFN-γ reactive band on immunoblot**

Whole cell lysate was mixed with sample loading buffer without the addition of denaturing or reducing agents. Samples were separated in polyacrylamide gels without addition of SDS after which they were electrotransferred to polyvinylidene difluoride (PVDF) membranes or stained with Coomassie Blue. Following the wash steps and blocking, membranes were incubated with 5 μg of IFN-γ overnight at 4°C. Membranes were then stained with anti-human IFN-γ antibody for 1 hour at room temperature followed by anti-mouse HRP for 1 hour. Membranes were developed with enhanced chemiluminescence reagent (Thermo Fisher) to detect immunoreactive bands. The corresponding bands on the Coomassie stained gel were excised and stored at -80°C until digestion.

**Protein identification by mass spectrometry**

The gel slices were then rinsed with 100 mM ammonium bicarbonate solution and transferred into a sterile Eppendorf®LoBind 1.5 ml microcentrifuge tube. A volume of 500 µl of acetonitrile (ACN) was added and the sample was incubated on ice, for 10 minutes. Following brief centrifugation, the ACN was removed and 100µl of 10 mM dithiothreitol (DTT) solution was added and incubated at 56°C for 30 min., removed and cooled to room temperature. Next, 500 µl of ACN was added and incubated on ice for a further 10 min after which the liquid solution was aspirated off, 100 µl of 55 mM iodoacetamide solution was added and the gel sample was incubated at room temperature for 30 min in the dark. Next, 500 µl of ACN was added and the sample was incubated on ice, for 10 min, the liquid solution was then aspirated and diluted trypsin (Promega, sequence grade) was added to each gel sample to a final concentration of 10 µg per 1 µl of ProteaseMAX™ (Promega) 1% trypsin enhancer solution. The sample was then mixed gently and incubated at 4°C for 2 hours. The samples were then incubated in the trypsin/ProteaseMAX™ solution at 37°C overnight. The resulting peptides were extracted by adding 400 µl of 5% formic acid/acetonitrile (1:2, v/v) solution to the sample followed by 15 min incubation at 37°C on a shaking heating block set at 450 rpm. The samples were then briefly centrifuged, the solution transferred to a sterile Eppendorf®LoBind microcentrifuge tube and the sample dried using a SpeedVac concentrator (Labconco, USA) set at 40°C. The dried, extracted peptides were reconstituted in 50 µl of 5% formic acid solution, and injected for nano-LC-MS/MS analysis. The peptide digests were analyzed using a shotgun analysis on a Thermo Q Exactive Orbitrap mass spectrometer coupled to a Dionex UltiMate 3000 UPLC system. The resultant Thermo RAW files were subjected to analysis using Thermo Proteome Discoverer 2.2, SEQUEST and the *Mtb* H37Rv protein FASTA file obtained from Uniprot. The peptide false discovery rate (FDR) set at <0.01 was used to get confident results.

**Enhancement of isoniazid killing of *Mtb* by IFN-γ**

Bacterial cultures were diluted in fresh 7H9 broth containing ADC. Subsequently, mid-log phase *Mtb* H37Rv was diluted to an $OD_{600}$ of 0.01 in 7H9 media (0.2% glycerol, 0.01% Tyloxapol, 10% ADC). These bacilli were aliquoted into inkwells and cultured at 37°C. Isoniazid was added at a concentration of 1 ug/mL; 100 ng/mL in the case for IFN-γ. 100 µl of the untreated and treated cultures was taken at indicated time points, serially diluted in phosphate buffered saline and plated onto Middlebrook 7H11 agar containing ADC.

**RNA sample preparation**

Bacteria were grown to an $OD_{600}$ of 0.6 and then pelleted and treated with RNAprotect Bacteria Reagent (Qiagen) for 15 min at room temperature. Samples were washed with PBS and resuspended in TRIzol (Life Technologies) and stored at -80°C. Thawed samples were transferred to Lysis Matrix B tubes containing 0.1-mm silica beads (Q-Biogene) and homogenized in a MagnaLyser instrument (Roche) at 7,000 rpm for 5 x 60 s with 3 min incubation on ice in between homogenizations. Samples were centrifuged for 1 min at 16,100 x g at 4°C, and the supernatant was transferred to a new Eppendorf tube. After phenol-chloroform extraction, the nucleic acids were precipitated with isopropanol, washed with 75% ethanol, air dried for 10 min, and finally resuspended in nuclease-free water. Genomic DNA was removed with RNase-Free DNase Set (Qiagen) and RNA was further purified on-column using RNeasy Mini Kit (Qiagen) and eluted in 50 µL of nuclease-free water. Purity and integrity were verified by Nanodrop (Thermo) and Bioanalyzer (Agilent) respectively.

**RNA sequencing analysis**

Paired-end sequence reads were quantified to transcript abundance using Kallisto with bias correction, and 50 bootstrap samples resulting in at least 35.9 million aligned reads per sample. Reads were mapped to *Mtb* H37Rv individual gene sequences database from Mycobrowser release 4. On average, the percentage of aligned reads was 89.4%. The transcript abundance was then summarized to gene level using Sleuth. Raw counts from RNA-sequencing were processed in Bioconductor package EdgeR, variance was estimated, and size factor normalized using Trimmed Mean of M-values (TMM). Genes were filtered using the filtered_p function of the genefilter package. All fit models included a term to model individual variation. For the identification of DEGs a group comparison was applied using experiment batch as covariant. Genes with a FDR-corrected p-value < 0.05 were identified as differentially expressed, resulting from a likelihood ratio test using a negative binomial generalized linear model fit. Top genes with a nominal p value < 0.01 were also considered for further validation. Pathway enrichment analysis was performed using the cellular overview tool available in the BioCyc database collection based on the *Mtb* H37Rv reference genome.

**Complementation of *mmpl10* in the mutant strain**

The *mmpl10* (*rv1183*) ORF was PCR amplified from *Mtb* genomic DNA using the primers (Thermo, USA) rv1183F(ATGTTCGAAGTGGTCGGCTGTTGGGTCGC), rv1183R(CACGTTAACCCGCCTTCGGCGGCTAAACA) and KOD Xtreme HotStart DNA polymerase (Roche). After PCR clean up, the *mmpl10* PCR product and pMV762 shuttle vector were digested with *Bst*BI and *Hpa*I (Thermo, USA), purified using agarose electrophoresis, and ligated with T4 DNA ligase (NEB) to produce pMV762::*mmpl10*. The pMV762::*mmpl10* complementation vector expressed *mmpl10* under the control the $hsp_{60}$ promoter and were electroporated using Gene Pulser Xcell (Biorad) into electrocompetent *Δmmpl10 Mtb* transposon mutant. Transformants were selected on 7H10 agar plates containing hygromycin (50 µg/mL).

**3D cell culture**

Microspheres were generated with an electrostatic generator (Nisco, Zurich, Switzerland) as described previously (*40*). Briefly, PBMC were infected overnight with *Mtb* in a 75 $cm^3$ flask, cells were detached, pelleted and mixed with 1.5% sterile alginate (Pronova UP MVG alginate, Nova Matrix, Norway) and 1 mg/mL collagen (Advanced BioMatrix, USA) at a final concentration of $5x10^6$ cells/ml. The cell-alginate suspension was injected into the bead generator where microspheres were formed in an ionotropic gelling bath of 100 mM $CaCl_2$ in HBSS. After washing twice with HBSS with $Ca^{2+}/Mg^{2+}$, microspheres were dispensed into eppendorfs and transferred in RPMI 1640 medium (Thermo Fisher) containing 25 µg/mL kanamycin, 1% ampicillin, 10% human AB serum and incubated at 37˚C, 5% $CO_2$. *Mtb* growth within microspheres was monitored longitudinally by luminescence (GloMax 20/20 Luminometer, Promega). In the case of IFN-γ supplementation, the media contained the indicated concentrations of cytokine. In the IFN-γ blockade experiment, the cell suspension was incubated with anti-IFN-γ antibody (Biolegend) for 1 hour at 4°C before addition to the alginate-collagen matrix. Time points described are days post infection.

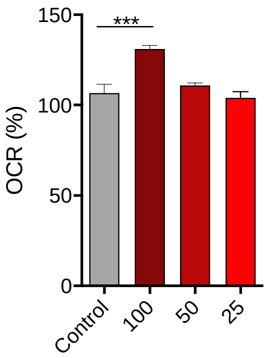


**Fig. S1. Murine recombinant IFN-γ increases *Mtb* OCR.**

The mouse is the most used animal model used for investigations into basic immunology during Mtb infection. Addition of recombinant murine IFN-γ to *Mtb* at indicated concentrations (ng/mL) induces dose-dependent increase in OCR similar to the effect seen with human IFN-γ. Data are shown as mean ± SEM and represent at a minimum two independent experiments. Tukey's correction multiple-comparison test was used for the statistical analysis.

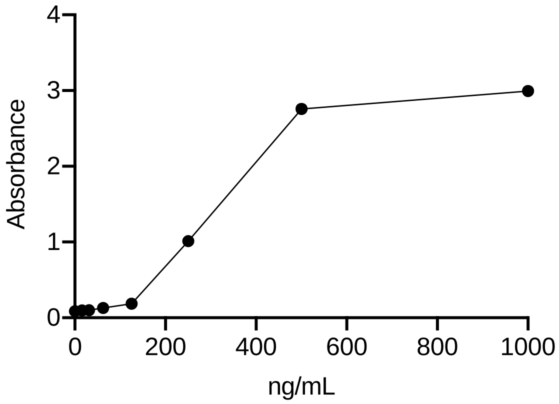


**Fig. S2. Binding of IFN-γ to *Mtb***

IFN-γ binds to formalin-fixed *Mtb* in a dose-dependent manner at indicated concentrations (ng/mL) as measured by ELISA. This assay complements the binding observed by flow cytometry and confocal microscopy. Data are shown as mean ± SEM and represent at a minimum two independent experiments.

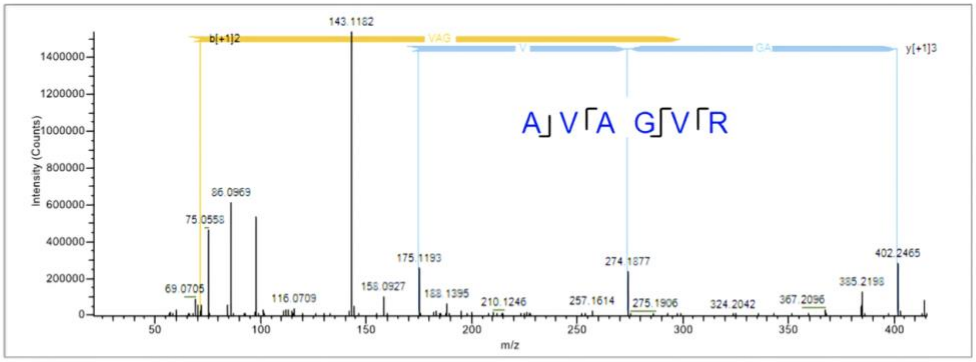


Fig. S3. **Ionic spectrum of unique MmpL10 fragment obtained by HCD fragmentation.**

Unique fragment spectrum to *Mtb* MmpL10 obtained by HCD fragmentation.

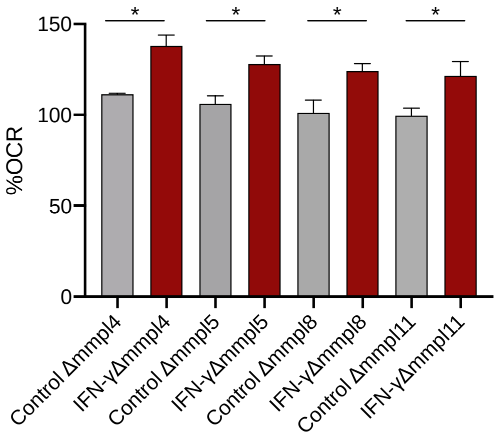


Fig. S4. **IFN-γ increases OCR in several *Mtb Δmmpls*.**

Human recombinant IFN-γ increases OCR in several *Mtb Δmmpls* to confirm this effect is mediated by MmpL10. Data are shown as mean ± SEM and represent at a minimum two independent experiments. Tukey's correction multiple-comparison test was used for the statistical analysis.

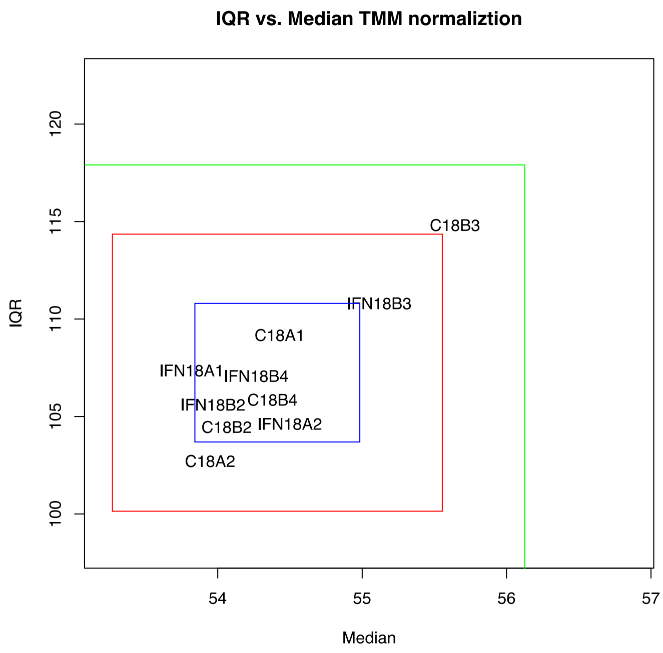


Fig. S5. **Interquartile range analysis of RNA sequencing**

Control samples denoted with C; IFN-γ-treated samples denoted by IFN. Five replicates per condition. Blue square indicates 1SD, red square 2SD and green square 3SD.

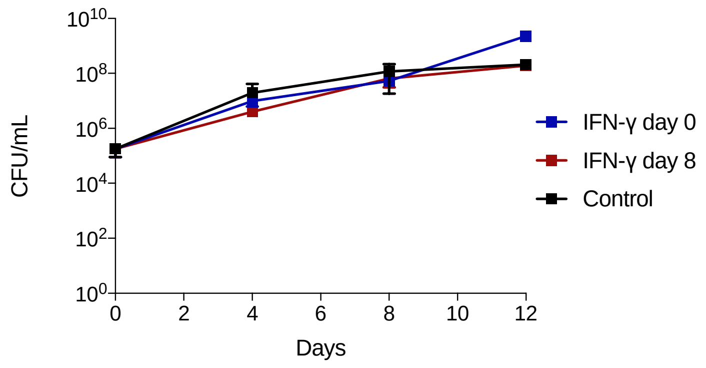


Fig. S6. **IFN-γ does not enhance growth of *Mtb* cultures**

Addition of human recombinant IFN-γ to *Mtb* only does not increase bacterial growth.

| Accession | Description | Coverage [%] | # Peptides | # PSMs | # Unique Peptic | # Protein Grour | # AAs | MW [kDa] | calc. pI | Score Sequest HT: Sequest HT |
|---|---|---|---|---|---|---|---|---|---|---|
| P9WPE7 | 60 kDa chaperonin 2 OS=Mycobacterium tuberculosis (strain ATCC 25618 / H37Rv) OX=83332 GN=groEL2 PE=1 SV=1 | 18 | 6 | 6 | 6 | 1 | 540 | 56,7 | 4,92 | 2.62 |
| P96400 | Probable conserved transmembrane protein OS=Mycobacterium tuberculosis (strain ATCC 25618 / H37Rv) OX=83332 GN=Rv0218 PE=4 SV=1 | 2 | 1 | 1 | 1 | 1 | 442 | 47,3 | 10,08 | 0.00 |
| P9WJU1 | Acyltrehalose exporter MmpL10 OS=Mycobacterium tuberculosis (strain ATCC 25618 / H37Rv) OX=83332 GN=mmpL10 PE=1 SV=1 | 1 | 1 | 4 | 1 | 1 | 1002 | 106,3 | 8,65 | 0.00 |
| O05790 | Possible phosphatase OS=Mycobacterium tuberculosis (strain ATCC 25618 / H37Rv) OX=83332 GN=Rv3113 PE=4 SV=1 | 8 | 1 | 1 | 1 | 1 | 222 | 24,8 | 5,67 | 0.00 |
| I6Y4V2 | Uncharacterized protein OS=Mycobacterium tuberculosis (strain ATCC 25618 / H37Rv) OX=83332 GN=Rv0804 PE=1 SV=1 | 14 | 1 | 1 | 1 | 1 | 209 | 21,6 | 12,10 | 0.00 |
| P9WHR3 | Carboxylesterase A OS=Mycobacterium tuberculosis (strain ATCC 25618 / H37Rv) OX=83332 GN=caeA PE=1 SV=1 | 6 | 1 | 1 | 1 | 1 | 520 | 55,9 | 6,19 | 0.00 |
| P9WKH1 | (2E,6E)-farnesyl diphosphate synthase OS=Mycobacterium tuberculosis (strain ATCC 25618 / H37Rv) OX=83332 GN=Rv3398c PE=1 SV=1 | 12 | 1 | 1 | 1 | 1 | 359 | 38,8 | 6,32 | 0.00 |
| P9WFQ3 | Nucleotide-binding protein Rv1421 OS=Mycobacterium tuberculosis (strain ATCC 25618 / H37Rv) OX=83332 GN=Rv1421 PE=1 SV=1 | 2 | 1 | 1 | 1 | 1 | 301 | 32,9 | 6,98 | 0.00 |
| I6X666 | Uncharacterized protein OS=Mycobacterium tuberculosis (strain ATCC 25618 / H37Rv) OX=83332 GN=Rv3076 PE=1 SV=1 | 5 | 1 | 2 | 1 | 1 | 158 | 17,1 | 9,95 | 0.00 |
| P9WN45 | 1,4-alpha-glucan branching enzyme GlgB OS=Mycobacterium tuberculosis (strain ATCC 25618 / H37Rv) OX=83332 GN=glgB PE=1 SV=1 | 3 | 1 | 1 | 1 | 1 | 731 | 81,7 | 5,73 | 0.00 |

**Table S1. Proteins in immunoreactive membrane bands corresponding to peptide fragments observed by mass spectrometry.**

Shotgun mass spectrometry detected several peptides in the immunoreactive band. Only MmpL10 is a membrane-associated protein

| ID | ORF size | POI | ORF description | Rv# | JHU_ID |
|---|---|---|---|---|---|
| HG1139 | 2904 | 311 | MmpL4 | Rv0450c | JHU0450c-311 |
| STN1424 | 2895 | 1540 | MmpL5 | Rv0676c | JHU0676c-1540 |
| JO1275 | 3270 | 114 | MmpL8 | Rv3823c | JHU3823c-114 |
| STN0573 | 3009 | 2396 | MmpL10 | Rv1183 | JHU1183-2396 |
| NA0234 | 2901 | 279 | MmpL11 | Rv0202c | JHU0202c-279 |

**Table S2. List of *Mtb Δmmpls***

List of all *Mtb Δmmpls* used in this study taken from JHU transposon mutant library

| Gene | C18A1 | IFN18A1 | C18A2 | IFN18A2 | C18B2 | IFN18B2 | C18B3 | IFN18B3 | C18B4 | IFN18B4 | logFC | logCPM | LR | PValue | FDR |
|---|---|---|---|---|---|---|---|---|---|---|---|---|---|---|---|
| Rv3561 | 2.195.142 | 198.632 | 3.008.154 | 2.551.917 | 3.000.314 | 2.501.671 | 7.962.967 | 6.115.893 | 4.266.514 | 3.819.546 | -0.24635 | 5.224.387 | 2.908.184 | 6.94E-08 | 0.000288 |
| MTB000021 | 1.504.634 | 1.773.455 | 1.203.052 | 1.337.287 | 1.167.094 | 1.271.669 | 1.003.249 | 1.316.903 | 1.780.877 | 1.932.131 | 0.203959 | 7.156.781 | 2.258.887 | 2.01E-06 | 0.002777 |
| Rv1686c | 9.040.289 | 7.875.697 | 1.647.472 | 1.370.094 | 1.581.817 | 1.238.235 | 5.571.255 | 3.837.003 | 1.668.652 | 1.565.082 | -0.31017 | 4.340.593 | 228.777 | 1.73E-06 | 0.002777 |
| Rv1687c | 1.151.263 | 9.963.231 | 2.442.948 | 1.856.618 | 2.312.168 | 1.774.187 | 8.532.087 | 6.234.833 | 2.457.172 | 2.583.628 | -0.28661 | 4.927.323 | 1.874.394 | 1.49E-05 | 0.015518 |
| Rv2620c | 1.548.337 | 1.745.938 | 3.447.759 | 2.259.258 | 3.273.737 | 2.251.336 | 1.218.903 | 1.052.381 | 4.132.368 | 3.347.537 | -0.3097 | 5.484.157 | 1.595.457 | 6.49E-05 | 0.053877 |
| Rv2347c | 1.532.075 | 171.611 | 8.478.865 | 9.440.587 | 9.055.374 | 1.011.146 | 9.184.787 | 1.196.144 | 1.593.022 | 1.654.109 | 0.182111 | 6.940.971 | 1.533.315 | 9.01E-05 | 0.062364 |
| Rv3559c | 1.198.712 | 1.173.447 | 1.331.375 | 1.062.522 | 1.223.981 | 1.117.268 | 2.060.118 | 1.629.477 | 1.675.196 | 1.388.079 | -0.22615 | 3.789.828 | 1.472.789 | 0.000124 | 0.073666 |
| Rv0393 | 3.789.283 | 3.185.071 | 4.224.395 | 428.737 | 4.621.408 | 3.998.642 | 3.967.373 | 338.741 | 3.060.127 | 2.599.154 | -0.17556 | 5.223.238 | 1.356.322 | 0.000231 | 0.090692 |
| Rv1953 | 212.272 | 2.530.344 | 2.009.623 | 2.852.033 | 203.691 | 2.721.764 | 2.963.183 | 4.329.414 | 2.879.243 | 3.012.162 | 0.379653 | 1.470.671 | 134.866 | 0.00024 | 0.090692 |
| Rv2618 | 2.816.974 | 2.941.525 | 3.747.109 | 31.391 | 3.633.407 | 3.099.787 | 9.783.208 | 7.314.807 | 364.486 | 3.310.273 | -0.20645 | 5.443.283 | 1.371.596 | 0.000213 | 0.090692 |
| Rv3562 | 1.892.967 | 1.698.494 | 1.948.916 | 1.638.521 | 1.789.178 | 1.634.739 | 4.465.941 | 3.480.183 | 24.539 | 2.347.623 | -0.20024 | 4.543.133 | 139.859 | 0.000184 | 0.090692 |
| Rv1845c | 1.617.263 | 1.446.724 | 1.361.938 | 1.341.201 | 1.441.802 | 1.272.509 | 1.969.341 | 1.642.799 | 1.365.023 | 1.293.987 | -0.14118 | 720.524 | 1.319.461 | 0.000281 | 0.097142 |
| Rv3341 | 5.728.846 | 5.544.617 | 6.518.714 | 5.845.736 | 6.538.298 | 5.307.441 | 1.298.392 | 9.308.241 | 6.877.464 | 6.744.758 | -0.20611 | 61.584 | 1.240.264 | 0.000429 | 0.136929 |
| Rv3190A | 5.886.177 | 6.373.305 | 4.354.183 | 4.524.107 | 4.398.625 | 4.736.206 | 3.226.577 | 4.291.354 | 4.780.197 | 5.412.575 | 0.170057 | 5.582.173 | 1.204.067 | 0.000521 | 0.154372 |
| Rv2621c | 1.184.478 | 1.370.182 | 2.815.147 | 2.114.978 | 282.58 | 2.013.266 | 7.503.203 | 5.438.648 | 2.675.733 | 233.427 | -0.27309 | 8.242.718 | 1.129.592 | 0.000777 | 0.215012 |

**Table S3. Extended list of differentially expressed *Mtb* genes**

All differentially regulated genes in Mtb after 18 hours IFN-γ incubation. Controls are denoted by C; IFN-γ are denoted by IFN.
